# The Role of the Eyes: Investigating Face Cognition Mechanisms Using Machine Learning and Partial Face Stimuli

**DOI:** 10.1101/2023.05.15.540886

**Authors:** Ingon Chanpornpakdi, Toshihisa Tanaka

**Author notes:** Corresponding author: Toshihisa Tanaka.

## Abstract

Face cognition plays a significant role in social interaction. The typical stimulus used to study face cognition mechanisms is a rapid serial visual presentation (RSVP). During the RSVP task, the brain response called event-related potential (ERP) is evoked when a person recognizes a target image. Many trials are required to average and obtain a clean ERP to interpret the cognitive mechanism behind the ERP response. However, increasing the trial number can cause fatigue and affect evoked ERP amplitude. This paper adopts a different perspective; machine learning might extract a meaningful cognitive result that reveals the face cognition mechanism without directly focusing on the characteristic of the ERP. We implemented an xDAWN covariance matrix method to enhance the data quality and a support vector machine classification model to predict the participant’s event of interest using ERP components evoked in the partial face cognition task. The effect of face components and the physical response was also investigated to explore the role of each component and find the possibility of reducing fatigue caused during the experiment. We found that the eyes were the most effective component. Similar statistical results were obtained from full face and partial face with eyes visible in both behavioral response and classification performance. From these results, the eye component could be the most crucial in face cognition. So, there could be some similarities in the face cognition mechanism of the full face and the partial face with eyes visible, which should be further investigated using ERP characteristics.

## I. INTRODUCTION

The SARS-CoV-2 pandemic brought not only a significant epidemiological experience for the people but also an unprecedented experience in terms of face recognition— wearing facial masks. Face cognition is a fundamental and essential skill in social interaction. When people interact, they first identify a person, then recognize social cues such as their facial position, emotional expressions, eye gaze, and physical changes to understand each other, decide how to make the next move, and establish a pleasant interaction. Covering part of the face forced the mechanism of face cognition to change from full to partial face. What are the psychophysiological effects of wearing a mask on face perception? This study addressed this question using electrophysiological measurements and machine learning.

Previous studies of partial face cognition reported that holistic face processing enhances the identification judgment and visual short- and long-term memory for face cognition [1]−[3]. Having part of the face covered weakens face cognition and matching performance abilities [1], [4]–[9]. Moreover, missing the eye component causes the loss of structural information. It delays the response time while missing the nose component leads to the loss of configural information about the face [10]. Despite many studies on partial face cognition, how people can recognize partial faces correctly is still in question.

This paper addressed the problem of partial face cognition with the rapid serial visual presentation (RSVP) paradigm, which is commonly used in visual perception studies [11]. It is a visual oddball task that presents a stream of target and non-target stimuli at an extremely high rate, during which the target is presented more infrequently. The stimuli used in the RSVP can be either static stimuli, such as images [12] or dynamic stimuli, such as videos [13]. A response to the RSVP can be captured by electroencephalography (EEG). During the RSVP task, an event-related potential (ERP) corresponding to cognition is induced when a participant perceives the target image. The induced ERP is a time-locked response associated with the stimulus onset [14], [15]. It is a temporal waveform with components named according to their polarity and latency. One of the most commonly used biomarkers for visual perception is P300, the positive peak induced 250–700 msec after stimulus onset responsible for the rare target perception [11]. The analysis of ERP (such as P300) typically involves the grand averaging procedure to the signal-to-noise ratio so that the obtained ERP has high consistency with less contamination of artifacts [16].

Many studies claim that averaging a few trials (14 to 20) could yield reliable ERP [17]–[19]. Still, it would preferably be averaged about 100 trials [20]; hence, the appropriate number of trials is still in question. The reason is that it depends on many factors, such as the types of task (auditory or visual task), the nature of the participants (age and health), the sample size, the focused ERP component, or even the noise environment of the experimental room [16], [17], [21], [22]. Although increasing the number of trials can improve data quality and improve the data quality, more extended experiments may cause fatigue which could lead to more noise contamination in the actual experiment [22].

Thus, in this paper, we applied a machine learning approach to decode the ERP evoked by target and non-target in RSVP tasks for further application with few trials [23], [24]. We hypothesized in this study 1) covering part of the face lowers the ability to recognize faces and results in lower classification accuracy, and 2) the button press and no-button press tasks would perform comparably so that we could reduce the workload by substituting the button press task with the no-button press task. We designed an experiment with full face and face with each component covered, aimed to compare the similarity of full face and partial face cognition using the machine learning model to classify the target and non-target images using evoked ERP features straightforwardly. An xDAWN covariance matrix, which maximizes the signal-to-signal-plus-noise (SSNR) [25], [26], with tangent space mapping [27] was used for feature extraction. Then, support vector machine (SVM) was used in the ERP classification of face cognition task [28], as the combination of xDAWN and SVM was suggested as a suitable model for face cognition in RSVP tasks [11]. Furthermore, we tried to reduce the fatigue caused during the experiment due to the physical response, which is a concern in the face cognition task using EEG. We also investigated how physical response (button press) affects the model’s accuracy.

## II. MATERIALS AND METHODS

### A. PARTICIPANTS

Eighteen participants (nine males and nine females with an average age of 26.05 *±*2.61 years and an age range of 22– 31 years) voluntarily participated in the experiment after providing written informed consent. The study was approved by the Research Ethics Committee of the Tokyo University of Agriculture and Technology (N02-14-E87 and N02-14-E92). All participants self-reported normal or corrected-to-normal vision and had no neurological disorders. None of the student participants were encouraged to participate in this experiment by their professors, nor did they obtain any credits for doing so.

### B. STIMULUS DESIGN

Colorful face images from a public dataset of the “Ethnic Origins of Beauty (Les origines de la beauté)” project by Ivanova N. (available at lesoriginesdelabeaute.com) [29] were used as an RSVP stimulus. The stimulus presentation was created using the Psychophysics Toolbox Version 3 [30] in MATLAB 2020a. Asian-like faces with black hair, dark eyes, and fair skin were selected due to the other-race effect or other-race bias, an observed phenomenon that people can learn and recognize faces of their own race faster and better than other races [31]–[34]. The ethnic group of the face images used in this experiment was Uyghurs, Han, Korean, Tuvan, and Buryat.

The images were resized to 710 ×555 pixels and trimmed to a visual angle of 7.4× 5.2 c/d so that only the face area was visible. Previous studies have shown that distinctive faces are easier to recognize than typical faces [35], [36]; therefore, one distinctive face was set as the target image based on the shape of the nose and mouth, which stood out from other faces. Since all images were unfamiliar to the participants, we allowed them to learn the target face by performing a training task.

The experiment was divided into two parts, a training task and a main task, which consisted of two tasks (a button press task and a no button press task). Each task consisted of seven blocks with seven face conditions: full face images condition (all parts of the face were exposed to the participants) and six partial face images conditions (the images with the eyes covered, images with the nose covered, images with the mouth covered, images with the eyes and nose covered, images with the eyes and mouth covered, and images with the nose and mouth covered). At the beginning of each task, the instructions with a full face target image were provided for the participant to remember.

### C. TRAINING TASK

The training task used three Asian-like faces: two non-target faces and one designated target face. Each face was randomly shown 10 times (30 trials in total), and each trial was presented until the participant responded by pressing a button. The odd-numbered participants were asked to respond by pressing the left button, and the even-numbered participants were asked to respond with the right button when the target face was shown to avoid a decision-making response bias [37]. Auditory feedback was provided for both correct and incorrect responses to help participants learn the target face more easily. In this task, both visual and audio stimuli were used simultaneously, the beeps at 800, 1,300, and 2,000 Hz were played for correct responses, and only the 800 Hz beep was played consecutively for incorrect responses. According to Armstrong and Issartel [38], there are discrepancies between the saliency of auditory stimuli when compared to visual stimuli since visual stimulus has more than one salient point. To overcome this biased, they suggested using audio with a frequency of 800 Hz. However, the audio stimulus was only used in the training task which there were no biosignal or behavioral responses recorded, the frequency used as audio stimuli might not have had a great impact on the EEG recording task.

### D. MAIN TASK

The stimulus presentation contained 120 trials with one target face trained in the training task and five non-target Asian-like faces that had never been used in the training task. Each face was randomly presented 20 times (0.16 probability rate of the target). The experimental flow of each block started with an instruction, followed by fixation for 500 msec to avoid attentional blinks, in which the second target image cannot be perceived up to 500 msec after the first target image is identified [39].

To investigate the effect of the button press response, two tasks (button press and no button press tasks) were performed. The stimulus presentation was set to 200 msec [40], [41]. However, the participant will unlikely respond by pressing the button within 200 msec of the presentation duration. Therefore, we have extended the presentation time to 1000 msec for the button press task. Consequently, the image stimuli were presented at 1,000 msec for the button press task and 200 msec for the no button press task, with 500 msec of fixation between each image. During the button press task, the participants were asked to respond when the target face was shown by clicking the left mouse. In addition, the participants were also asked to count the number of the target faces they saw and fill in the number at the end of each block for both tasks 1. In the no-button press task, the participants were asked to count the number of the target faces seen in the same manner as the button press task. EEG, eye movements, button press action (only for the button press task), and the number of target faces recognized were recorded.

### E. DATA ACQUISITION

EEG was recorded at a sampling frequency of 2,048 Hz using Polybench software (Twente Medical Systems International B.V., TMSi, Oldenzaal, Netherlands). Participants wore a 64-electrode EEG gel head cap, based on the 10-10 international system (Fp1, Fpz, Fp2, AF7, AF3, AF4, AF8, F7, F5, F3, F1, Fz, F2, F4, F6, F8, FT7, FC5, FC3, FC1, FCz, FC2, FC4, FC6, FT8, M1, T3, C5, C3, C1, Cz, C2, C4, C6, T4, M2, TP7, CP5, CP3, CP1, CPz, CP2, CP4, CP6, TP8, T5, P5, P3, P1, Pz, P2, P4, P6, T6, PO7, PO5, PO3, POz, PO4, PO6, PO8, O1, Oz, and O2). A disposable ground electrode was placed on the participant’s left wrist to reduce artifact interference compared to the EEG ground electrode, as suggested by the TMSi manual. As a reference, the average of the input EEG signals was amplified using a 72-channel Refa Amplifier (TMSi, Oldenzaal, Netherlands). Eye movements were measured simultaneously to detect eye blinks. They were recorded with two channels, vertical and horizontal, using a pair of microelectrodes (bipolar, TMSi, Oldenzaal, Netherlands). The vertical eye movement channel was placed above the right eye with a reference electrode on the left ear lobe. The horizontal eye movement channel was placed near the outer canthus of the right eye with a reference electrode on the right ear lobe. Each participant placed their head on chin support and sat 60 cm from a VIEWpixx display (120-Hz refresh rate, 1, 920 ×1, 200 pixels, Vpixx Technologies, Saint Bruno, Canada) in a soundproof room.

### F. DATA ANALYSIS

#### 1) Pre-processing

The recorded EEG was analyzed using MNE package 1.3.0 [42] in Python 3.5.8. The data were first re-referenced using an average of the EEG around the earlobe electrodes (M1 and M2) and filtered with one to 30 Hz FIR bandpass filter. The artifacts such as eye blink were removed using an extended infomax independent component analysis (ICA) algorithm on the continuous data. The 64 electrodes were grouped into 12 groups based on the brain lobe: frontal, parietal, occipital, and temporal area (Figure 2, Table 1) [43] and combined into one electrode by means. The obtained EEG data (120 trials) were sliced into 120 epochs from 100 msec before stimulus onset to 1,000 msec. To confirm the quality of data, we have calculated the signal-to-noise ratio on the grand average of each participant was calculated for each condition (both button press and no button press) by dividing the P300 peak amplitude by the standard deviation of the amplitude across the pre-stimulus interval (*-*100 to 0 msec) [44]–[46])

**TABLE 1.**
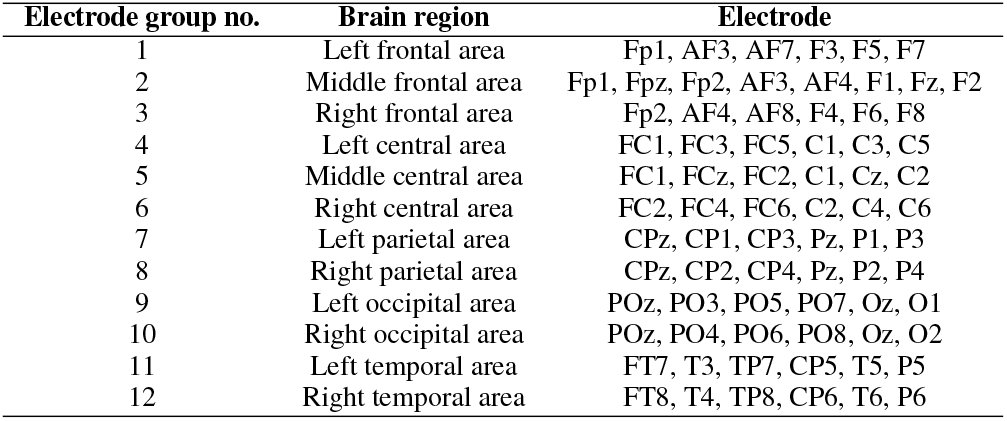
The brain region and EEG electrode position correspond to the brain lobe

**FIGURE 1.**
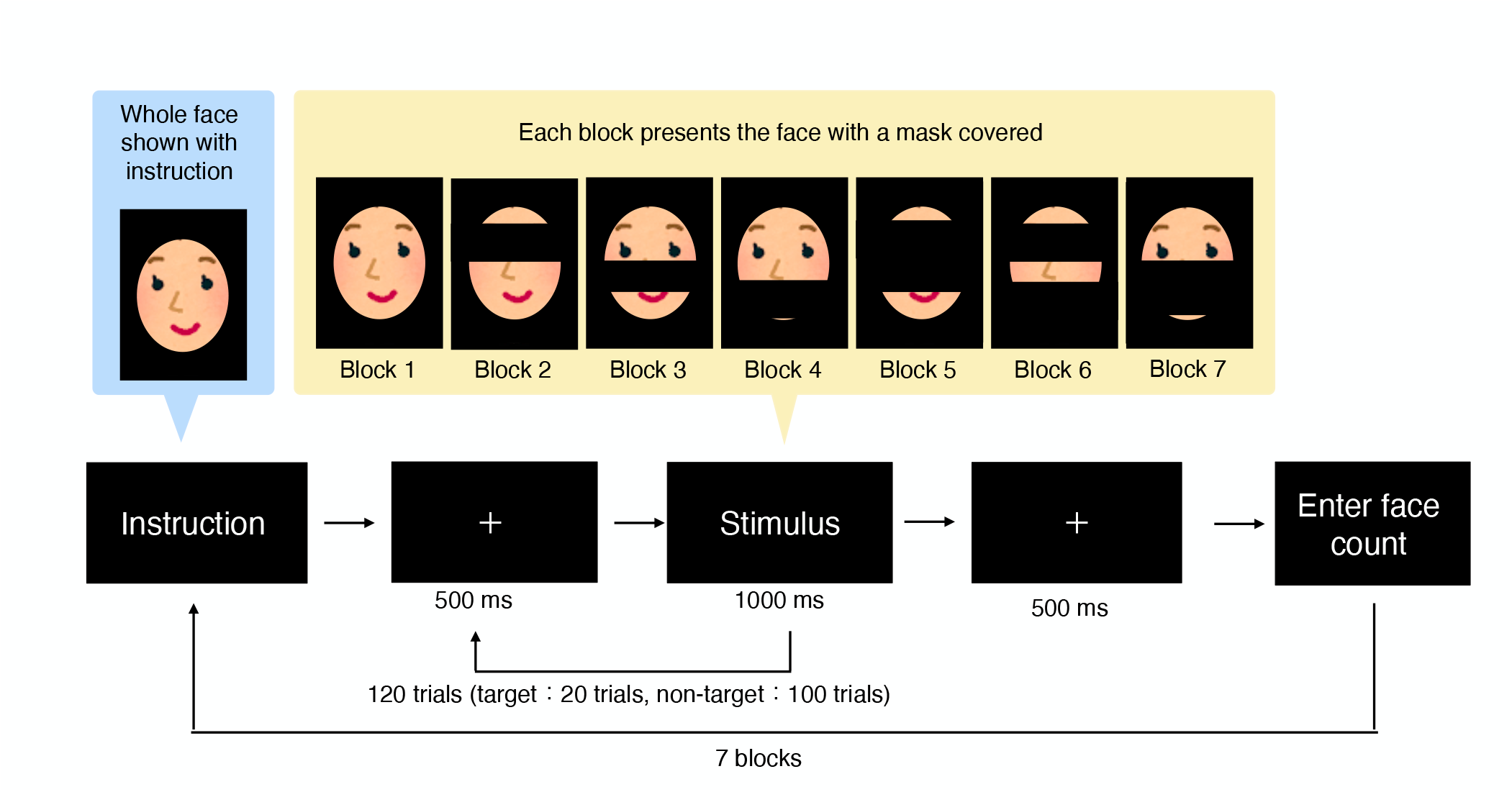
Experimental flow for the main tasks (The face images used in the real experiment were the dataset from Les origines de la beauté project)

**FIGURE 2.**
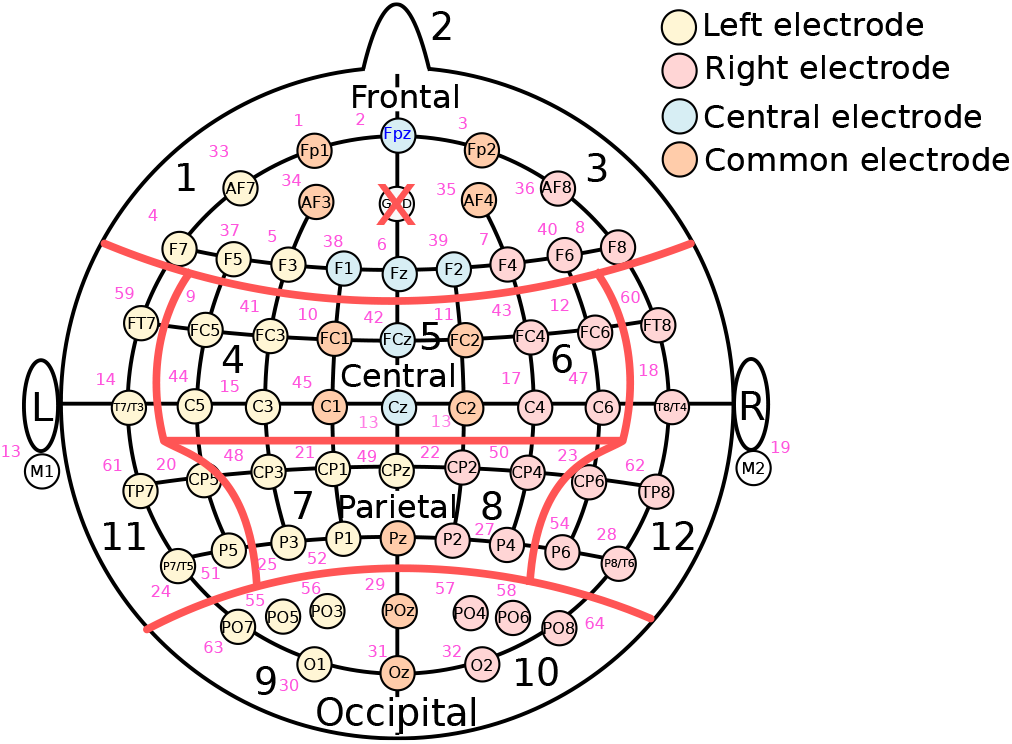
Electrode group with yellow representing the left hemisphere, red representing the right hemisphere, blue representing the center of the EEG electrode position, and orange representing the common electrodes in their corresponding groups.

#### 2) Behavioral Response

The button press error count was calculated by counting the number of times that participants missed the task or press the button incorrectly. If the target face was shown and participants did not press the button, it was considered an error count for the target. On the other hand, if the participants pressed the button when the target face was not shown, it was considered an error count for the non-target. Two-way repeated measures analysis of variance (ANOVA) with Greenhouse-Geisser correction in multiple comparisons was used to calculate the button press error rate of 18 participants, defined as

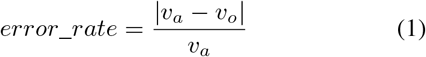

where *v*_*a*_ and *v*_*o*_ denote the ideal value (*v*_*a*_ = 20) and observed value, respectively. Eq. (1) yields the error rate of the button press response for each face and the error rate of the target face count of each task.

The two factors used in ANOVA analysis were the type of faces and face cognition conditions. The hypotheses were 1) all the face types had an equal mean of error rate, 2) all the face cognition conditions had an equal mean of error rate, and 3) the error rate of face types and face cognition conditions were independent of each other. The target face error count on both button press and no button press tasks were calculated by subtracting the face count number from the correct number of 20 target faces and using the absolute value of that face error count to find the total face error count among 18 participants. However, the target face error count’s results were in doubt because the count on the correct target image could not be clarified. There are chances that the participants could have counted on the non-target and coincidentally got the count correctly. For that reason, we cannot take this result into account and did not perform the statistical test for the face count error rate.

#### 3) Machine Learning-Based Analysis

The whole epoch of the pre-processed data was used in both the training and testing models. The data contained 20 trials of target images and 100 trials of non-target images, with 12 electrode groups and 2,254 samples each. Since the data were imbalanced, the non-target data were divided into five sets by selecting four trials from each non-target face without replacement. This resulted in 20 trials for each set, with each non-target face event equally distributed. With this method, 40 trials for each set of data were achieved when combining the non-target data with the target data. The obtained data were randomized and inputted into a supervised learning model to classify the brain response patterns of the target and non-target images. In addition, the extraction of features using the xDAWN covariance matrix with tangent space mapping [27].

The xDAWN is the spatial filter which is given as

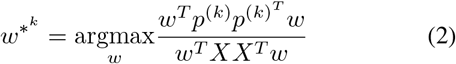

where *w* ∈ ℝ^*E*×1^, *E* is the number of electrodes, *k* denotes the class index, *P* ∈ ℝ^*E*×*T*^ represents the grand average of the *E* electrodes ti *T* time samples and *X* ∈ ℝ^*E*×*T*^ is the signal including two classes: 0 and 1 [25], [26]. The matrix 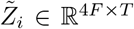 of filtered evoke potential of both classes and the filtered data were used to calculate the covariance matrix ∑_*i*_ ∈ ℝ^4*F* ×4*F*^ where *F* is the number of xDAWN filters. The matrix 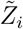 is defined as

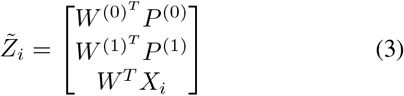

where *W* ^(*k*)^ ∈ ℝ^*E*×*F*^ is the selected filter of class *k* and *i* is the index of a trial from *N*_*k*_ trials, in which 1 ≤*i* ≤ *N*_*k*_ [27].

The extracted features were classified by a linear SVM model to identify the target face. The accuracy of each model was calculated by averaging five accuracies obtained from each set of data. Two-way repeated measures ANOVA with Greenhouse-Geisser correction was applied to our accuracy results to examine our hypotheses that 1) all the face condition groups have equal mean accuracies, 2) all the button press conditions have equal mean accuracies, and 3) the factors are independent, or the interaction effect of the face cognition conditions and button press conditions does not exist. The two factors in our ANOVA test were seven face cognition conditions (full face and six partial face conditions) and button press conditions. The dependent variable was the number of participants, which was 18 in this study. After identifying the conditions that were significantly different, pairwise t-tests were performed to explore the relationships that showed significant differences using Benjamini/Hochberg (non-negative) correction.

## III. RESULTS

### A. BEHAVIORAL RESPONSE

Figures 3 and 4 show the total error count of the button press response and the target face count of 18 participants, respectively (Supplementary Tables I and II for error rate details for each subject). From Figure 3, we can see that the button press error count was highest when the eyes and mouth component of the face was covered and second highest when the eyes and mouth were covered. On the other hand, the button press error count was similar in full face cognition and partial face cognition with the eyes component presented.

**FIGURE 3.**
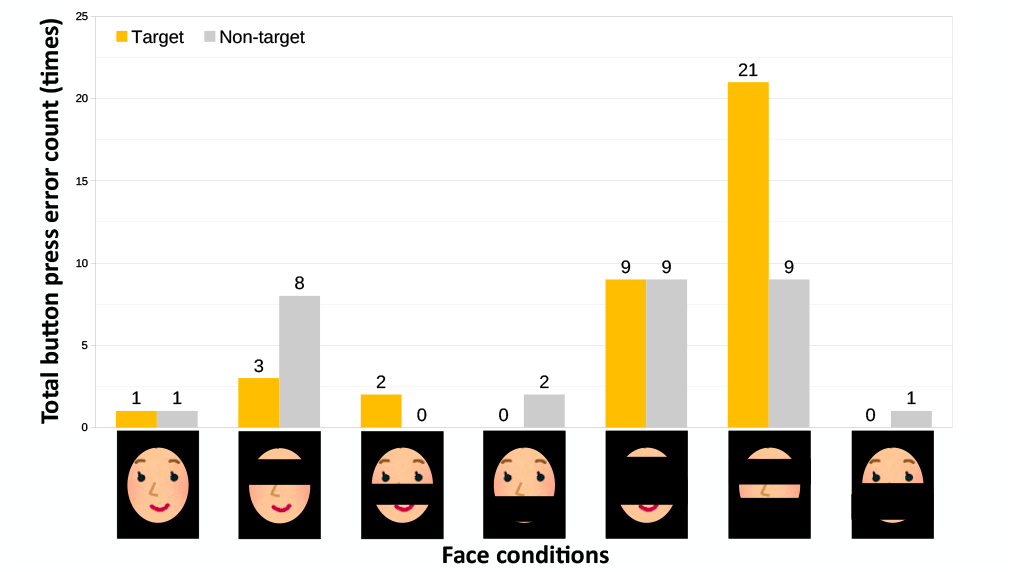
Total button press error count of 18 participants in the button press task, where yellow and gray colors represent the button press error count when the target face image and non-target face image are shown, respectively. The total number of errors for the target face image was 360, and that for the non-target face image was 1,800.

**FIGURE 4.**
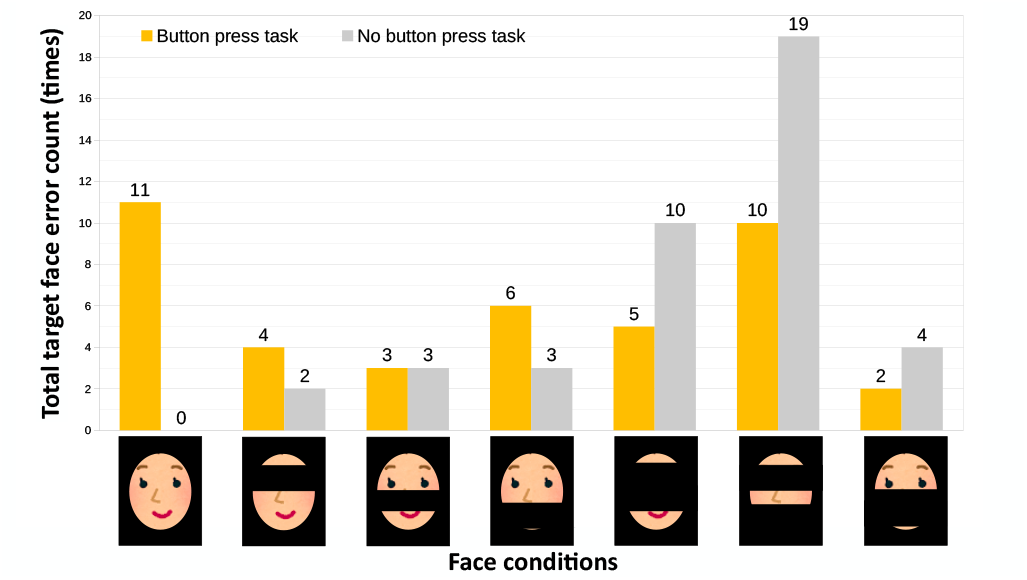
Total number of target face error counts of 18 participants in both button press and no button press tasks, where yellow and grey colors represent the error count in the button press and no button press tasks, respectively. The total number of errors for both the button press and no button press tasks was 360

Significant differences were also found between the face cognition condition (*F* (6, 102) = 9.681,*p <* 0.001, *η*_*p*_^2^ = 0.209), but could not find the difference in the type of faces that participants made mistakes while pressing the button; missing the button press on the target face and pressing the button during non-target face presentation (*F* (1, 17) = 0.500,*p >* 0.05, *η*_*p*_^2^ = 0.002). No interaction between the face cognition condition and the type of face where the error occurred was found (*F* (6, 102) = 3.239,*p <* 0.05, *η*_*p*_^2^ = 0.058) (see Table 2). A pairwise t-test shows that the button press error rate during the face with eyes and mouth covered is significantly different from all other conditions except the face with eyes and nose covered; full face (*p <* 0.001), the face with eyes covered (*p <* 0.05), the face with nose covered (*p<* 0.001), the face with mouth covered (*p<* 0.001), the face with nose and mouth covered (*p <* 0.001). In addition, the pairwise t-test also shows the button press error rate of the condition when the face with eyes and nose covered has significant differences between the full face (*p<* 0.05), the face with nose covered (*p<* 0.05), and the face with nose and mouth covered (*p <* 0.05) (see the t-values and effect sizes in Supplementary Table III).

**TABLE 2.** The two-way repeated measures ANOVA result of button press error rate in button press task.

| Source | SS | df1 | df2 | MS | F | p-unc | p-GG-corr | $\eta_p^2$ | eps |
| --- | --- | --- | --- | --- | --- | --- | --- | --- | --- |
| Condition | 0.051 | 6 | 102 | 0.0085 | 9.691 | 1.980E-08 | 0.000 | 0.209 | 0.423 |
| Face type | 0.000 | 1 | 17 | 0.0004 | 0.500 | 0.489 | 0.489 | 0.002 | 1.000 |
| Condition * Face type | 0.012 | 6 | 102 | 0.0020 | 3.239 | 0.006 | 0.045 | 0.058 | 0.371 |

Similarly, we could see from Figure 4 that the target face count rate was higher when the eye component was covered, especially when either nose or mouth was also covered in the no-button press task. However, the target face error count was relatively high during full face cognition in the button press task which was opposed to the button press error count.

### B. CLASSIFICATION PERFORMANCE

Figure 5 shows the example of ERP obtained from the full face and the partial face cognition in the button press task. The ERP was obtained by grand averaging the EEG data of 20 trials across 18 participants. We could see that the P300 evoked by the target image was relatively larger when compared with that of the non-target images.

**FIGURE 5.**
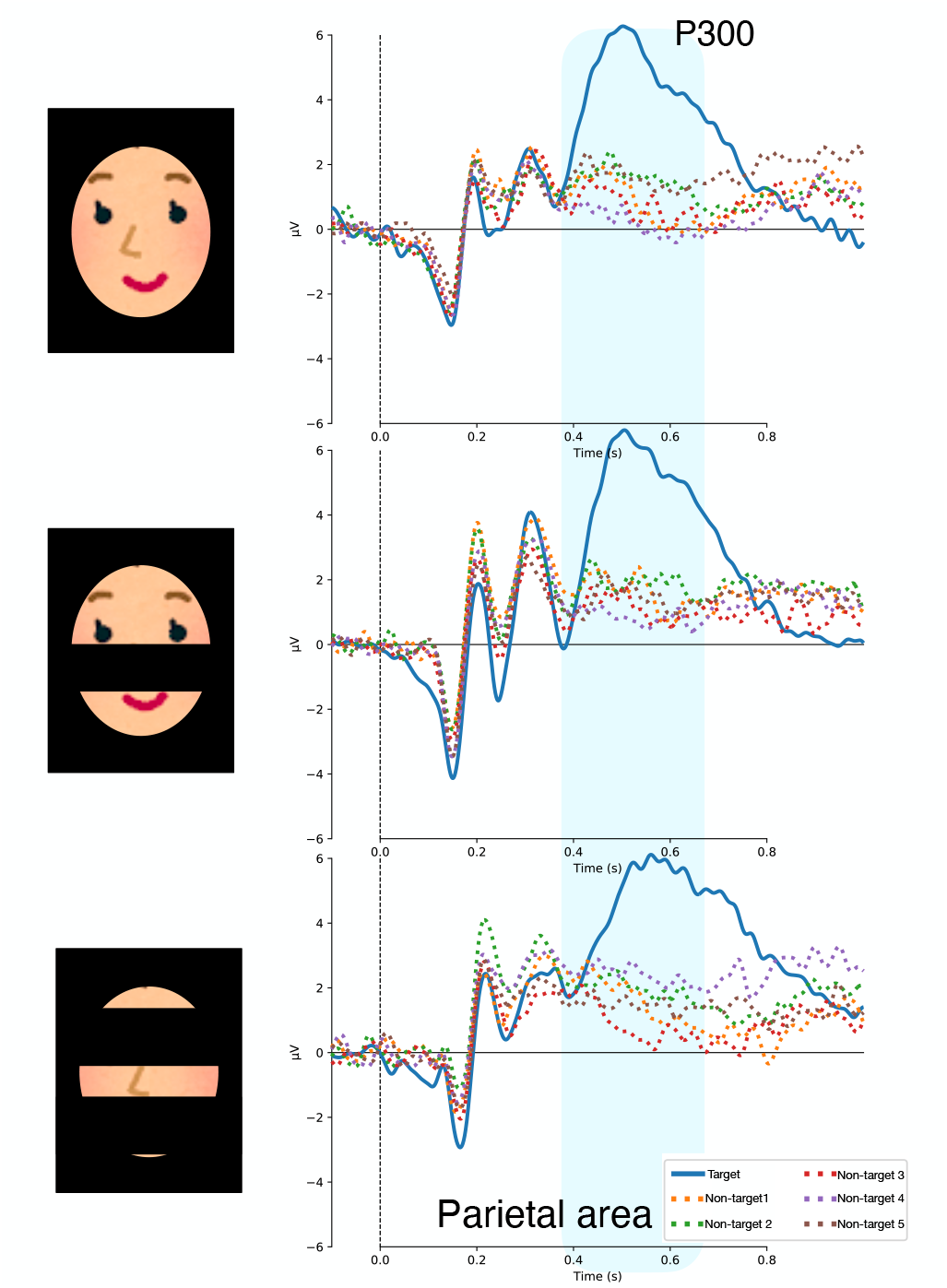
P300 evoked by the target and non-target images during full face and partial face cognition condition in button press task at the parietal area. The conditions shown were a face with a nose covered as an example of a partial face with one face component covered and a face with eyes and mouth covered as an example of a partial face with two face components covered. The ERP was obtained by grand averaging across 18 participants and the P300 component was highlighted in blue color. The blue line represents ERPs evoked by the target image and the dotted line represents the ERPs evoked by non-target images.

We inputted the processed EEG data through the xDAWN filter and linear SVM model for target and non-target classification. The results of the classification model are shown in Figure 6. It represents the average accuracy across the five datasets extracted by balancing the ERPs evoked by target and non-target faces during full and partial face cognition in both button press and no-button press tasks. The accuracy was calculated by averaging the accuracy obtained from 18 participants. We could see that the accuracy of the button press task was almost equal to the accuracy of the no-button press task except when the face with nose covered, face with mouth covered and face with nose and mouth covered. In the mentioned three conditions, the accuracy during the button press task was slightly higher than the no-button press task. The highest accuracy of 0.860 when the participants recognized the face with the nose and mouth covered during the button press task. In contrast, the highest accuracy for the no-button press task was 0.862 obtained when the participant recognized the full face.

**FIGURE 6.**
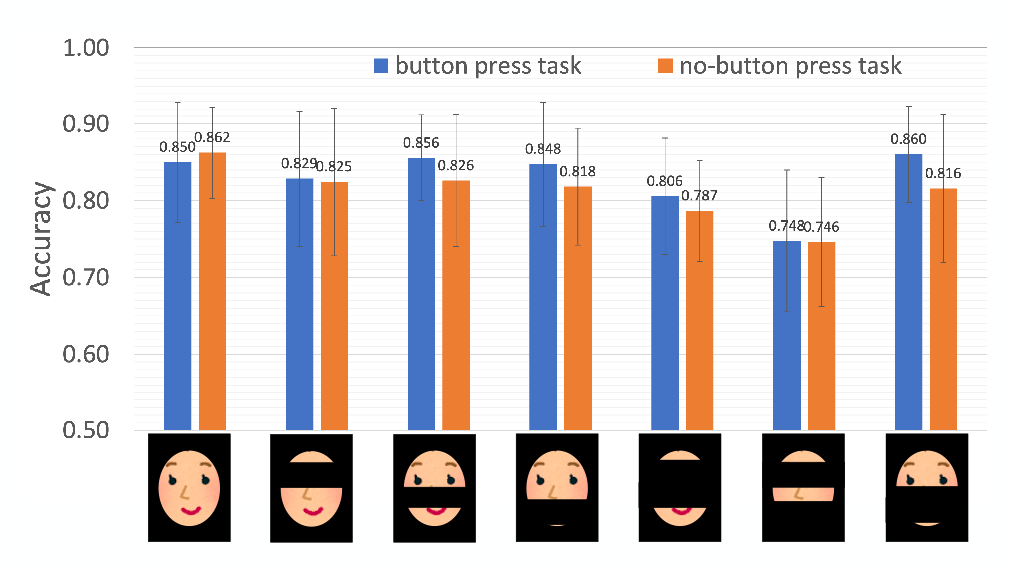
The accuracy of participant dependent cross-validation in target face classification using xDAWN covariance matrix, tangent space mapping, and linear SVM model where blue color represents button press task and orange color represents no-button press task. The error bar denotes the standard deviation around the mean.

Moreover, when comparing the accuracy of the face cognition condition with eyes being invisible including, the differences in accuracy changed between the two physical response conditions were unremarkable (Figure 6). The accuracy also became lowest when the eyes and mouth were covered in both button press and no-button press tasks.

The two-way repeated measures ANOVA test of the ERP classification accuracy results were shown in Table 3. We could see that the accuracy in ERP classification significantly affected only the face cognition condition groups (*F* (6, 102) = 11.139,*p <* 0.001, *η*_*p*_^2^ = 0.164), but no significant difference in the button press conditions (*F* (1, 17) = 1.409,*p* = 0.252, *η*_*p*_^2^ = 0.011), and the interaction between the face cognition conditions and button press conditions (*F* (6, 102) = 1.201,*p* = 0.317, *η*_*p*_^2^ = 0.014) were found.

**TABLE 3.** The two-way repeated measures ANOVA result of target and non-target classification accuracy for full and partial face cognition in button press and no-button press task.

| Source | SS | df1 | df2 | MS | F | p-unc | p-GG-corr | $\eta_p^2$ | eps |
| --- | --- | --- | --- | --- | --- | --- | --- | --- | --- |
| Condition | 0.295 | 6 | 102 | 0.049 | 11.139 | 1.576e-09 | 4.529e-07 | 0.164 | 0.675 |
| Button | 0.0174 | 1 | 17 | 0.017 | 1.409 | 0.252 | 0.252 | 0.011 | 1.000 |
| Condition * Button | 0.021 | 6 | 102 | 0.004 | 1.201 | 0.312 | 0.317 | 0.014 | 0.758 |

The pairwise t-test shows the significant differences between the accuracy of the face with eyes and mouth covered condition and the other face condition; full face (*p <* 0.005), the face with eyes covered (*p <* 0.001), the face with nose covered (*p <* 0.005), the face with mouth covered (*p<* 0.001), the face with eyes and nose covered (*p<* 0.01), and the face with nose and mouth covered (*p <* 0.005). Furthermore, significant differences between the face with eyes and nose covered condition and the following four face conditions were observed; the full face (*p <* 0.01), the face with nose covered (*p <* 0.05), and the face with mouth covered (*p <* 0.05) (see the t-values and effect sizes in Supplementary Tables IV).

## IV. DISCUSSION

In this study, we performed classification using P300 data in the supervised model to explore the role of the face components in face cognition. We found that the accuracy dropped when the eyes and nose were covered and became lowest among all the conditions when the eyes and mouth were covered. We also investigate our hypothesis that the no physical response (no button press) task would lower the fatigue caused during the cognition task. We found that the ERP evoked in the button press task provided better accuracy than the no button press task in partial face cognition. However, no significant differences were observed between the accuracy of the button press and the no-button press task.

### A. BEHAVIORAL RESPONSE

The behavior responses of the button press, and target face count error rate showed comparable results that the error became higher when the eyes and nose or mouth were covered. However, no statistical test was performed for the target face count error rate due to the reliability of the data. On the contrary, a significant difference was found in the button error rate in the condition when the eyes and another component were missing compared to the other face conditions.

When the eye components were hidden together with another face component, the error rate of both the button press response and the target face count increased and was markedly higher than the task with only more components hidden. These results indicated that many participants made more mistakes when eye components were invisible and made the face cognition task become harder when the other component of the face was covered along with the eyes and suggesting that eye components might be crucial in face cognition. We also observed that the error rate of the face count was not associated with that of the button response as we saw that the full face target image count error rate was higher than when the eyes were covered in the button press task result. Regardless of the higher target face count error rate, participants were able to respond to the target image by pressing the button correctly. Some participants also reported that they were too focused on the button press task and might have miscounted the target face number, which caused a higher error rate in the target face count. However, as we mentioned in the Methods section the result of face count might not be accurate and reliable enough to make a conclusion. Only the number of times the participants counted the face was recorded. The issue was that if the participants counted on the wrong face but got the number of counts correct, it would be recorded as a correct task, and without clarification, such a flaw could lead to the wrong result if the face count is also considered. For that reason, we neglected the results of the face count error rate.

Our result obtained in this behavior responses showed that the button press error rate was similar in the case of full face and partial face with the eyes visible. These results implied that there is a possibility that people can recognize a partial face with the eyes at a similar level as in full face cognition. However, many studies reported that covering parts of the face such as having a mask on the face reduces the ability of face perception especially for face identification [5], [6] and emotional categorization [7]–[9]. One reason why our result of the partial face with the nose and mouth covered (equivalent to wearing a mask) had a similar level of button press task error might be that we trained the participants to learn the face prior to the experiment and they performed the same target face cognition task for 7 blocks. The participants might become more familiar with both the target face and the partial face cognition task as they performed the experiment and make the cognition of the partial face with the nose and mouth covered, which is the last block to perform, became easier than it could be.

### B. TARGET CLASSIFICATION PERFORMANCE

Although the number of trials was only 20 per image type, the reason that the ERP data yielded high accuracy in target image classification is that we used the xDAWN covariance matrix for feature extraction. It is calculated with the matrix of both filtered average data or evoked and filtered data which improves the SNRR [27]. As a result, we found that the accuracy decreased only when the eyes and nose were covered and became lowest when the eyes and mouth were covered. We also observed that the highest accuracy in the button press task was when the participants recognized the face with a nose and mouth covered whereas the highest accuracy in the no-button press task was during the full face cognition. This result can be interpreted that the presence of eyes plays a crucial role in partial face cognition [4], [47] which supports our behavioral response finding.

In addition, the accuracy in the button press task and no-button press task was almost the same in all the face cognition conditions except in the partial face cognition with the eyes visible. In partial face cognition with the eyes visible (the face with nose covered, the face with mouth covered, and the face with nose and mouth covered), the accuracy in the button press task was higher than the no-button press task but no significant differences were observed between the accuracy of the two tasks. Gerson et al. [48] studied the effect of button press and no button press on the surveillance RSVP task by presenting the images for 100 msec and found similar results to our finding that there were no significant differences in the button press and no-button press task performance. Although the button press response might not affect the classification performance, we should further investigate the ERP characteristic changes in both button response tasks to achieve a stronger conclusion.

In the comparison of behavioral response and classification performance statistical results, the two-way repeated measures ANOVA results in both behavioral response and classification performance show significant differences in the face cognition condition. In particular, we observed that the face with eyes and nose covered and the face with eyes and mouth covered have significant differences in accuracy when compared with the full face, the face with nose covered and the face with mouth covered in both behavioral responses and classification performance results. From these results, we could assume that the full face, the face with the nose covered, and the face with the mouth covered might have a similar cognitive response pattern, having eyes as the most important component.

### C. EFFECT ON EXPERIMENTAL DESIGN

We only recorded the button press accuracy to confirm the behavioral responses. Nevertheless, to understand the brain motor activity of the button press mechanism, the reaction time of the button press is also an important key that could provide a fruitful meaning to our results. In addition, the presentation time for the button press task was set to 1,000 msec, whereas in the no button press task, it was set to 200 msec. The significant difference in the presentation time can cause a better perception of the button press task due to a longer time to expose to the stimulus and process the presented information of the image. Consequently, the target count number might be inaccurate due to shorter or longer memory maintenance periods and the combination of all these factors. This difference in time could also affect the accuracy as previous studies [49] found that the classification performance of the RSVP task improves when presented with the images for 100–200 msec compared with that of 50 msec presentation time.

To overcome such a problem, we should also investigate a similar presentation period to diminish any inconsistency in our experiment and investigate the effect of workload in button press tasks. It was suggested that 500 msec could be the most suitable for the RSVP face cognition task [11] Presenting the face image stimulus for 500 msec might also yield a better brain response in the no-button press task and reduce the accuracy-workload trade-off in the classification model.

### D. LIMITATION AND FUTURE WORK

Several factors such as the length of the experiment [50] can also affect the quality of EEG when the experiment is run for a long time. These factors can lower the ERP level, resulting in lower accuracy, and hence, they also need to be investigated along with the button press task. In the next stage, we would like to examine the mechanism of partial face cognition by exploring the characteristics of ERP: amplitude, and latency, with a more significant trial number of the task to compare our classification results so that this study can reveal more details on how to face components affect the brain response during the face cognition and the role of button press task.

## ACKNOWLEDGMENT

This work was supported in part by JST Moonshot R&D Grant Number JPMJMS2237.

