## Supplementary Table for "The Role of the Eyes: Investigating Face Cognition Mechanisms Using Machine Learning and Partial Face Stimuli"

Supplementary Table I. The button response error rate of 18 participants

| **Cond.** | **Full face** | | | | | | **Covered eyes** | | | | | | **Covered nose** | | | | | | **Covered mouth** | | | | | | **Covered eyes and nose** | | | | | | **Covered eyes and mouth** | | | | | | **Covered nose and mouth** | | | | | |
| --- | --- | --- | --- | --- | --- | --- | --- | --- | --- | --- | --- | --- | --- | --- | --- | --- | --- | --- | --- | --- | --- | --- | --- | --- | --- | --- | --- | --- | --- | --- | --- | --- | --- | --- | --- | --- | --- | --- | --- | --- | --- | --- |
| **Face** | **T** | **1** | **2** | **3** | **4** | **5** | **T** | **1** | **2** | **3** | **4** | **5** | **T** | **1** | **2** | **3** | **4** | **5** | **T** | **1** | **2** | **3** | **4** | **5** | **T** | **1** | **2** | **3** | **4** | **5** | **T** | **1** | **2** | **3** | **4** | **5** | **T** | **1** | **2** | **3** | **4** | **5** |
| P1 | **0.05** | **0** | **0** | **0** | **0** | **0** | **0.05** | **0** | **0** | **0** | **0** | **0** | **0.05** | **0** | **0** | **0** | **0** | **0** | **0** | **0** | **0** | **0** | **0** | **0** | **0.1** | **0.05** | **0** | **0** | **0** | **0** | **0.05** | **0** | **0** | **0** | **0** | **0** | **0** | **0** | **0** | **0** | **0** | **0** |
| P2 | **0** | **0** | **0** | **0** | **0** | **0** | **0** | **0** | **0** | **0** | **0** | **0** | **0** | **0** | **0** | **0** | **0** | **0** | **0** | **0** | **0** | **0** | **0** | **0** | **0.05** | **0** | **0** | **0** | **0** | **0.1** | **0.05** | **0** | **0** | **0** | **0** | **0.05** | **0** | **0** | **0** | **0** | **0** | **0** |
| P3 | **0** | **0** | **0** | **0** | **0** | **0** | **0** | **0.05** | **0** | **0** | **0** | **0** | **0** | **0** | **0** | **0** | **0** | **0** | **0** | **0** | **0** | **0** | **0** | **0** | **0** | **0** | **0** | **0.05** | **0** | **0** | **0.10** | **0** | **0** | **0** | **0** | **0.05** | **0** | **0** | **0.05** | **0** | **0** | **0** |
| P4 | **0** | **0** | **0** | **0** | **0** | **0** | **0** | **0** | **0** | **0** | **0** | **0** | **0** | **0** | **0** | **0** | **0** | **0** | **0** | **0** | **0** | **0** | **0** | **0** | **0.05** | **0** | **0** | **0** | **0** | **0** | **0.05** | **0** | **0.05** | **0** | **0** | **0** | **0** | **0** | **0** | **0** | **0** | **0** |
| P5 | **0** | **0** | **0** | **0** | **0** | **0** | **0** | **0** | **0** | **0** | **0** | **0** | **0** | **0** | **0** | **0** | **0** | **0** | **0** | **0** | **0** | **0** | **0** | **0** | **0** | **0** | **0** | **0** | **0** | **0** | **0** | **0** | **0** | **0** | **0** | **0** | **0** | **0** | **0** | **0** | **0** | **0** |
| P6 | **0** | **0** | **0** | **0** | **0** | **0** | **0.1** | **0** | **0** | **0** | **0** | **0.05** | **0** | **0** | **0** | **0** | **0** | **0** | **0** | **0** | **0** | **0** | **0** | **0** | **0** | **0** | **0** | **0** | **0** | **0** | **0.05** | **0** | **0** | **0** | **0** | **0** | **0** | **0** | **0** | **0** | **0** | **0** |
| P7 | **0** | **0** | **0** | **0** | **0** | **0** | **0** | **0.05** | **0** | **0** | **0** | **0** | **0** | **0** | **0** | **0** | **0** | **0** | **0** | **0** | **0** | **0** | **0** | **0** | **0** | **0** | **0** | **0** | **0** | **0.05** | **0.05** | **0** | **0** | **0** | **0** | **0** | **0** | **0** | **0** | **0** | **0** | **0** |
| P8 | **0** | **0** | **0** | **0** | **0** | **0** | **0** | **0** | **0** | **0** | **0** | **0** | **0.05** | **0** | **0** | **0** | **0** | **0** | **0** | **0** | **0** | **0** | **0** | **0** | **0** | **0** | **0** | **0** | **0** | **0** | **0.1** | **0** | **0** | **0** | **0** | **0** | **0** | **0** | **0** | **0** | **0** | **0** |
| P9 | **0** | **0** | **0** | **0** | **0** | **0** | **0** | **0** | **0** | **0** | **0** | **0** | **0** | **0** | **0** | **0** | **0** | **0** | **0** | **0** | **0** | **0** | **0** | **0** | **0.05** | **0** | **0** | **0** | **0** | **0.1** | **0.05** | **0** | **0** | **0** | **0** | **0.05** | **0** | **0** | **0** | **0** | **0** | **0** |
| P10 | **0** | **0** | **0** | **0** | **0** | **0** | **0** | **0** | **0** | **0** | **0** | **0.15** | **0** | **0** | **0** | **0** | **0** | **0** | **0** | **0** | **0** | **0** | **0** | **0** | **0.15** | **0** | **0** | **0** | **0** | **0.05** | **0.25** | **0** | **0.05** | **0** | **0** | **0** | **0** | **0** | **0** | **0** | **0** | **0** |
| P11 | **0** | **0** | **0** | **0** | **0.05** | **0** | **0** | **0** | **0** | **0** | **0** | **0** | **0** | **0** | **0** | **0** | **0** | **0** | **0** | **0** | **0** | **0** | **0** | **0.1** | **0** | **0** | **0** | **0** | **0** | **0** | **0** | **0** | **0** | **0** | **0** | **0** | **0** | **0** | **0** | **0** | **0** | **0** |
| P12 | **0** | **0** | **0** | **0** | **0** | **0** | **0** | **0** | **0** | **0** | **0** | **0** | **0** | **0** | **0** | **0** | **0** | **0** | **0** | **0** | **0** | **0** | **0** | **0** | **0** | **0** | **0** | **0** | **0** | **0** | **0** | **0** | **0** | **0** | **0** | **0** | **0** | **0** | **0** | **0** | **0** | **0** |
| P13 | **0** | **0** | **0** | **0** | **0** | **0** | **0** | **0.05** | **0** | **0** | **0** | **0** | **0** | **0** | **0** | **0** | **0** | **0** | **0** | **0** | **0** | **0** | **0** | **0** | **0** | **0** | **0** | **0** | **0** | **0.05** | **0.05** | **0** | **0** | **0** | **0** | **0** | **0** | **0** | **0** | **0** | **0** | **0** |
| P14 | **0** | **0** | **0** | **0** | **0** | **0** | **0** | **0** | **0** | **0** | **0** | **0** | **0** | **0** | **0** | **0** | **0** | **0** | **0** | **0** | **0** | **0** | **0** | **0** | **0** | **0** | **0** | **0** | **0** | **0** | **0.1** | **0** | **0** | **0** | **0** | **0** | **0** | **0** | **0** | **0** | **0** | **0** |
| P15 | **0** | **0** | **0** | **0** | **0** | **0** | **0** | **0** | **0** | **0** | **0** | **0** | **0** | **0** | **0** | **0** | **0** | **0** | **0** | **0** | **0** | **0** | **0** | **0** | **0** | **0** | **0** | **0** | **0** | **0** | **0** | **0** | **0** | **0** | **0** | **0** | **0** | **0** | **0** | **0** | **0** | **0** |
| P16 | **0** | **0** | **0** | **0** | **0** | **0** | **0** | **0** | **0** | **0** | **0** | **0** | **0** | **0** | **0** | **0** | **0** | **0** | **0** | **0** | **0** | **0** | **0** | **0** | **0** | **0** | **0** | **0** | **0** | **0** | **0.05** | **0** | **0** | **0** | **0.05** | **0** | **0** | **0** | **0** | **0** | **0** | **0** |
| P17 | **0** | **0** | **0** | **0** | **0** | **0** | **0** | **0.05** | **0** | **0** | **0** | **0** | **0** | **0** | **0** | **0** | **0** | **0** | **0** | **0** | **0** | **0** | **0** | **0** | **0** | **0** | **0** | **0** | **0** | **0** | **0.05** | **0** | **0** | **0** | **0** | **0** | **0** | **0** | **0** | **0** | **0** | **0** |
| P18 | **0** | **0** | **0** | **0** | **0** | **0** | **0** | **0** | **0** | **0** | **0** | **0** | **0** | **0** | **0** | **0** | **0** | **0** | **0** | **0** | **0** | **0** | **0** | **0** | **0.05** | **0** | **0** | **0** | **0** | **0** | **0.05** | **0** | **0.15** | **0** | **0** | **0** | **0** | **0** | **0** | **0** | **0** | **0** |

Note. T: target face image, 1, 2, 3, 4, 5: five different non-target face images, P: participant

Supplementary Table II. The target face count error rate of 18 participants in button press and no button press task.

| **Cond.** | **Full face** | | **Covered eyes** | | **Covered nose** | | **Covered mouth** | | **Covered eyes and nose** | | **Covered eyes and mouth** | | **Covered nose and mouth** | |
| --- | --- | --- | --- | --- | --- | --- | --- | --- | --- | --- | --- | --- | --- | --- |
| **Task** | **B** | **NB** | **B** | **NB** | **B** | **NB** | **B** | **NB** | **B** | **NB** | **B** | **NB** | **B** | **NB** |
| P1 | 0.1 | 0 | 0.05 | 0.05 | 0.05 | 0.05 | 0.05 | 0.05 | 0.05 | 0.05 | 0.05 | 0.05 | 0.05 | 0.05 |
| P2 | 0 | 0 | 0 | 0 | 0 | 0 | 0 | 0 | 0 | 0 | 0 | 0 | 0 | 0 |
| P3 | 0 | 0 | 0 | 0 | 0 | 0 | 0.05 | 0 | 0.05 | 0 | 0 | 0.2 | 0.05 | 0 |
| P4 | 0 | 0 | 0 | 0 | 0 | 0 | 0 | 0 | 0 | 0 | 0.05 | 0 | 0 | 0 |
| P5 | 0 | 0 | 0 | 0 | 0 | 0 | 0 | 0 | 0 | 0 | 0 | 0 | 0 | 0 |
| P6 | 0 | 0 | 0 | 0 | 0 | 0 | 0 | 0 | 0 | 0 | 0 | 0 | 0 | 0.05 |
| P7 | 0 | 0 | 0 | 0 | 0 | 0 | 0 | 0 | 0 | 0.05 | 0 | 0 | 0 | 0 |
| P8 | 0 | 0 | 0 | 0 | 0 | 0 | 0 | 0 | 0 | 0 | 0.25 | 0.25 | 0 | 0 |
| P9 | 0 | 0 | 0 | 0 | 0 | 0 | 0 | 0 | 0 | 0 | 0 | 0 | 0 | 0 |
| P10 | 0 | 0 | 0.1 | 0 | 0 | 0 | 0.05 | 0 | 0.1 | 0.1 | 0.1 | 0.15 | 0 | 0 |
| P11 | 0 | 0 | 0 | 0 | 0 | 0 | 0 | 0 | 0 | 0 | 0 | 0 | 0 | 0 |
| P12 | 0 | 0 | 0 | 0 | 0 | 0 | 0 | 0 | 0 | 0 | 0 | 0 | 0 | 0 |
| P13 | 0.05 | 0 | 0.05 | 0 | 0 | 0 | 0.05 | 0.05 | 0.05 | 0 | 0 | 0 | 0 | 0 |
| P14 | 0 | 0 | 0 | 0 | 0 | 0.05 | 0 | 0 | 0 | 0.20 | 0 | 0.1 | 0 | 0 |
| P15 | 0 | 0 | 0 | 0 | 0 | 0 | 0 | 0 | 0 | 0 | 0 | 0 | 0 | 0 |
| P16 | 0 | 0 | 0 | 0 | 0 | 0 | 0 | 0 | 0 | 0 | 0 | 0 | 0 | 0 |
| P17 | 0 | 0 | 0 | 0.05 | 0 | 0.05 | 0 | 0.05 | 0 | 0 | 0.05 | 0.05 | 0 | 0.1 |
| P18 | 0.4 | 0 | 0 | 0 | 0.1 | 0 | 0.1 | 0 | 0 | 0.1 | 0 | 0.15 | 0 | 0 |

Note. B: button press task, NB: no button press task, P: participant

Supplementary Table III. The pairwise t-tests result of button press error rate. (p < 0.05 highlighted in red)

| Contrast | Condition | A | B | Paired | Parametric | T | Degree of freedom | Tail of the test | Uncorrected p-values | Corrected p-values | P-values correction method | Bayes Factor | Hedges effect size |
| --- | --- | --- | --- | --- | --- | --- | --- | --- | --- | --- | --- | --- | --- |
| Condition | - | eyes | eyes and mouth | TRUE | TRUE | -3.04E+00 | 17 | two-sided | 0.00743 | 0.029103 | fdr_bh | 6.77 | -8.02E-01 |
| Condition | - | eyes | eyes and nose | TRUE | TRUE | -1.20E+00 | 17 | two-sided | 0.247586 | 0.34662 | fdr_bh | 0.451 | -3.26E-01 |
| Condition | - | eyes | full | TRUE | TRUE | 2.03E+00 | 17 | two-sided | 0.057917 | 0.093558 | fdr_bh | 1.287 | 6.71E-01 |
| Condition | - | eyes | mouth | TRUE | TRUE | 1.84E+00 | 17 | two-sided | 0.082698 | 0.124047 | fdr_bh | 0.981 | 6.37E-01 |
| Condition | - | eyes | nose | TRUE | TRUE | 2.03E+00 | 17 | two-sided | 0.057917 | 0.093558 | fdr_bh | 1.287 | 6.71E-01 |
| Condition | - | eyes | nose and mouth | TRUE | TRUE | 2.40E+00 | 17 | two-sided | 0.028326 | 0.059484 | fdr_bh | 2.26 | 7.63E-01 |
| Condition | - | eyes and mouth | eyes and nose | TRUE | TRUE | 2.13E+00 | 17 | two-sided | 0.048156 | 0.091935 | fdr_bh | 1.485 | 4.54E-01 |
| Condition | - | eyes and mouth | full | TRUE | TRUE | 3.99E+00 | 17 | two-sided | 0.000942 | 0.006592 | fdr_bh | 39.387 | 1.37E+00 |
| Condition | - | eyes and mouth | mouth | TRUE | TRUE | 3.83E+00 | 17 | two-sided | 0.001337 | 0.007019 | fdr_bh | 29.091 | 1.34E+00 |
| Condition | - | eyes and mouth | nose | TRUE | TRUE | 4.18E+00 | 17 | two-sided | 0.000633 | 0.006592 | fdr_bh | 55.652 | 1.37E+00 |
| Condition | - | eyes and mouth | nose and mouth | TRUE | TRUE | 4.56E+00 | 17 | two-sided | 0.000281 | 0.005892 | fdr_bh | 113.348 | 1.44E+00 |
| Condition | - | eyes and nose | full | TRUE | TRUE | 2.85E+00 | 17 | two-sided | 0.011087 | 0.029103 | fdr_bh | 4.853 | 8.99E-01 |
| Condition | - | eyes and nose | mouth | TRUE | TRUE | 2.53E+00 | 17 | two-sided | 0.021587 | 0.050369 | fdr_bh | 2.813 | 8.72E-01 |
| Condition | - | eyes and nose | nose | TRUE | TRUE | 2.85E+00 | 17 | two-sided | 0.011087 | 0.029103 | fdr_bh | 4.853 | 8.99E-01 |
| Condition | - | eyes and nose | nose and mouth | TRUE | TRUE | 2.97E+00 | 17 | two-sided | 0.008588 | 0.029103 | fdr_bh | 5.999 | 9.68E-01 |
| Condition | - | full | mouth | TRUE | TRUE | 0.00E+00 | 17 | two-sided | 1 | 1 | fdr_bh | 0.243 | 0.00E+00 |
| Condition | - | full | nose | TRUE | TRUE | 0.00E+00 | 17 | two-sided | 1 | 1 | fdr_bh | 0.243 | 0.00E+00 |
| Condition | - | full | nose and mouth | TRUE | TRUE | 5.66E-01 | 17 | two-sided | 0.578555 | 0.714686 | fdr_bh | 0.28 | 1.92E-01 |
| Condition | - | mouth | nose | TRUE | TRUE | 0.00E+00 | 17 | two-sided | 1 | 1 | fdr_bh | 0.243 | 0.00E+00 |
| Condition | - | mouth | nose and mouth | TRUE | TRUE | 4.37E-01 | 17 | two-sided | 0.667577 | 0.77884 | fdr_bh | 0.265 | 1.46E-01 |
| Condition | - | nose | nose and mouth | TRUE | TRUE | 5.66E-01 | 17 | two-sided | 0.578555 | 0.714686 | fdr_bh | 0.28 | 1.92E-01 |
| Face type | - | non-target | target | TRUE | TRUE | -7.07E-01 | 17 | two-sided | 0.489081 | NaN | NaN | 0.303 | -1.78E-01 |
| Condition * face type | eyes | non-target | target | TRUE | TRUE | 1.32E+00 | 17 | two-sided | 0.205291 | 0.463866 | fdr_bh | 0.511 | 4.10E-01 |
| Condition * face type | eyes and mouth | non-target | target | TRUE | TRUE | -2.29E+00 | 17 | two-sided | 0.035284 | 0.246989 | fdr_bh | 1.897 | -6.62E-01 |
| Condition * face type | eyes and nose | non-target | target | TRUE | TRUE | 7.79E-17 | 17 | two-sided | 1 | 1 | fdr_bh | 0.243 | 8.63E-17 |
| Condition * face type | full | non-target | target | TRUE | TRUE | 0.00E+00 | 17 | two-sided | 1 | 1 | fdr_bh | 0.243 | 0.00E+00 |
| Condition * face type | mouth | non-target | target | TRUE | TRUE | 1.00E+00 | 17 | two-sided | 0.331333 | 0.463866 | fdr_bh | 0.376 | 3.26E-01 |
| Condition * face type | nose | non-target | target | TRUE | TRUE | -1.46E+00 | 17 | two-sided | 0.163139 | 0.463866 | fdr_bh | 0.598 | -4.75E-01 |
| Condition * face type | nose and mouth | non-target | target | TRUE | TRUE | 1.00E+00 | 17 | two-sided | 0.331333 | 0.463866 | fdr_bh | 0.376 | 3.26E-01 |

Supplementary Table IV. The pairwise t-tests result of the accuracies. (p < 0.05 highlighted in red)

| **Contrast** | **Condition** | **A** | **B** | **Paired** | **Parametric** | **T** | **Degree of freedom** | **Tail of the test** | **Uncorrected p-values** | **Corrected p-values** | **P-values correction method** | **Bayes Factor** | **Hedges effect size** |
| --- | --- | --- | --- | --- | --- | --- | --- | --- | --- | --- | --- | --- | --- |
| **Condition** | - | eyes | eyes and mouth | TRUE | TRUE | 4.334206 | 17 | two-sided | 0.000451 | 0.001892 | fdr_bh | 74.835 | 1.073135 |
| **Condition** | - | eyes | eyes and nose | TRUE | TRUE | 1.756694 | 17 | two-sided | 0.096966 | 0.16969 | fdr_bh | 0.871 | 0.41676 |
| **Condition** | - | eyes | full | TRUE | TRUE | -2.465238 | 17 | two-sided | 0.024637 | 0.051737 | fdr_bh | 2.528 | -0.423582 |
| **Condition** | - | eyes | mouth | TRUE | TRUE | -0.458414 | 17 | two-sided | 0.652461 | 0.721141 | fdr_bh | 0.267 | -0.086418 |
| **Condition** | - | eyes | nose | TRUE | TRUE | -1.046642 | 17 | two-sided | 0.309919 | 0.406768 | fdr_bh | 0.392 | -0.193617 |
| **Condition** | - | eyes | nose and mouth | TRUE | TRUE | -0.725863 | 17 | two-sided | 0.477798 | 0.58077 | fdr_bh | 0.307 | -0.154396 |
| **Condition** | - | eyes and mouth | eyes and nose | TRUE | TRUE | -3.488015 | 17 | two-sided | 0.002816 | 0.008449 | fdr_bh | 15.357 | -0.764712 |
| **Condition** | - | eyes and mouth | full | TRUE | TRUE | -6.694283 | 17 | two-sided | 0.000004 | 0.000079 | fdr_bh | 5338.609 | -1.799755 |
| **Condition** | - | eyes and mouth | mouth | TRUE | TRUE | -5.428572 | 17 | two-sided | 0.000045 | 0.000474 | fdr_bh | 571.941 | -1.330234 |
| **Condition** | - | eyes and mouth | nose | TRUE | TRUE | -4.587563 | 17 | two-sided | 0.000262 | 0.001716 | fdr_bh | 120.426 | -1.452055 |
| **Condition** | - | eyes and mouth | nose and mouth | TRUE | TRUE | -4.483917 | 17 | two-sided | 0.000327 | 0.001716 | fdr_bh | 99.146 | -1.37996 |
| **Condition** | - | eyes and nose | full | TRUE | TRUE | -3.672873 | 17 | two-sided | 0.001886 | 0.0066 | fdr_bh | 21.642 | -1.012498 |
| **Condition** | - | eyes and nose | mouth | TRUE | TRUE | -2.687787 | 17 | two-sided | 0.015567 | 0.036322 | fdr_bh | 3.671 | -0.580917 |
| **Condition** | - | eyes and nose | nose | TRUE | TRUE | -2.786733 | 17 | two-sided | 0.012654 | 0.033216 | fdr_bh | 4.352 | -0.705128 |
| **Condition** | - | eyes and nose | nose and mouth | TRUE | TRUE | -2.221065 | 17 | two-sided | 0.04022 | 0.076784 | fdr_bh | 1.711 | -0.64785 |
| **Condition** | - | full | mouth | TRUE | TRUE | 1.529668 | 17 | two-sided | 0.144493 | 0.233412 | fdr_bh | 0.652 | 0.390037 |
| **Condition** | - | full | nose | TRUE | TRUE | 1.197574 | 17 | two-sided | 0.247515 | 0.346521 | fdr_bh | 0.452 | 0.258339 |
| **Condition** | - | full | nose and mouth | TRUE | TRUE | 1.219285 | 17 | two-sided | 0.239384 | 0.346521 | fdr_bh | 0.461 | 0.295617 |
| **Condition** | - | mouth | nose | TRUE | TRUE | -0.69278 | 17 | two-sided | 0.497803 | 0.58077 | fdr_bh | 0.301 | -0.123504 |
| **Condition** | - | mouth | nose and mouth | TRUE | TRUE | -0.363054 | 17 | two-sided | 0.721039 | 0.757091 | fdr_bh | 0.258 | -0.079566 |
| **Condition** | - | nose | nose and mouth | TRUE | TRUE | 0.208686 | 17 | two-sided | 0.837173 | 0.837173 | fdr_bh | 0.248 | 0.041189 |
| **Button** | - | button | no button | TRUE | TRUE | 1.18686 | 17 | two-sided | 0.251604 | NaN | NaN | 0.447 | 0.286809 |
| **Condition * Button** | eyes | button | no button | TRUE | TRUE | 0.185751 | 17 | two-sided | 0.854838 | 0.957782 | fdr_bh | 0.247 | 0.042605 |
| **Condition * Button** | eyes and mouth | button | no button | TRUE | TRUE | 0.053724 | 17 | two-sided | 0.957782 | 0.957782 | fdr_bh | 0.243 | 0.016957 |
| **Condition * Button** | eyes and nose | button | no button | TRUE | TRUE | 1.159185 | 17 | two-sided | 0.262404 | 0.459208 | fdr_bh | 0.435 | 0.26226 |
| **Condition * Button** | full | button | no button | TRUE | TRUE | -0.603553 | 17 | two-sided | 0.554109 | 0.775753 | fdr_bh | 0.286 | -0.174604 |
| **Condition * Button** | mouth | button | no button | TRUE | TRUE | 1.307685 | 17 | two-sided | 0.208389 | 0.459208 | fdr_bh | 0.506 | 0.366399 |
| **Condition * Button** | nose | button | no button | TRUE | TRUE | 1.71063 | 17 | two-sided | 0.105334 | 0.368671 | fdr_bh | 0.82 | 0.399515 |
| **Condition * Button** | nose and mouth | button | no button | TRUE | TRUE | 1.935267 | 17 | two-sided | 0.069772 | 0.368671 | fdr_bh | 1.116 | 0.534157 |
